# Rapid Pliocene Diversification of Modern Kangaroos

**DOI:** 10.1101/323717

**Authors:** Aidan M. C. Couzens, Gavin J. Prideaux

## Abstract

Differentiating between ancient and rapidly-evolved clades is critical for understanding impacts of environmental change on biodiversity. Australia possesses many aridity-adapted lineages, the origins of which have been linked by molecular evidence to late Miocene drying. Using dental macrowear and molar crown-height measurements spanning the past 25 million years, we show that the most iconic of Australia’s terrestrial mammals, ‘true’ kangaroos and wallabies (Macropodini), diversified in response to Pliocene grassland emergence. In contrast, low-crowned short-faced kangaroos radiated into browsing niches as the late Cenozoic became more arid, contradicting the view that this was a period of global decline among browsers. Our results link warm intervals with bursts of diversification and undermine arguments attributing Pleistocene megafaunal extinction to aridity-forced dietary change.

## Main Text

Adaptive radiation within novel ecological contexts is believed to underpin the diversification of many animal groups (*1*). However, the rapid divergence of lineages (*2*) can makes it challenging to accurately date their origins from molecular or morphological data (*3*). Often the result is a dichotomy between so-called short-and long-fuse diversification models, which can imply starkly different scenarios for how environmental change shapes biological diversity (*4*).

During the last 15 million years (Myr), Australia has undergone a large-scale environmental shift from a mesic to largely arid continent (*5*). This environmental shift has been closely linked with the evolution of a diverse, aridity-adapted biota (*6*), which includes such iconic members as the kangaroos and wallabies (Macropodidae), the most diverse marsupial herbivores ever to evolve. But, the timing and drivers of macropodid evolution have been difficult to resolve due to a patchy fossil record and imprecise divergence dating (*7*). Most phylogenetic analyses find that the Macropodini, the clade composed of species with grass-based diets (>70% of extant macropodoid diversity) was well underway during the arid late Miocene (c. 12–7 Myr ago) (*7–,!9*), 3–8 Myr before Australian grasslands emerged (*5*).

On the northern continents late Neogene grassland expansion ushered in the diversification of herbivores with high-crowned molar teeth, most notably ungulates (*10, 11*). Their success has been linked to the improved resistance of their dentitions to the elevated rates of dental wear characteristic of grazing diets (*12*). Similarly, macropodin grass consumers have higher-crowned molars than their browsing and fungivorous counterparts (*13*). The origin of high-crowned dentitions near the base of Macropodini (*7*) suggests that increased grass exploitation may have been a key factor in macropodin success. However, molar crown-height data has never been used to infer precisely when kangaroos became grazers or how this event was linked to environmental change.

To examine the role of dietary change in kangaroo adaptive radiation we measured molar macrowear levels (figs. S1,2) and crown height for >3,000 macropodoid specimens from the modern fauna and c. 100 fossil assemblages of late Oligocene to Holocene age (*14*). This enabled us to track diet and trait evolution in parallel. Our analysis reveals that macropodin kangaroos underwent a rapid burst of morphological diversification as grassland biomes took hold across Pliocene Australia.

Ancestral-state reconstruction shows that the low-crowned dentitions of balbarines and stem macropodids were ancestral for Macropodoidea (figs. 1A, 2A). Both groups have low crown heights (fig 1A) and limited macrowear disparity (fig 1B) suggesting either dietary overlap or that other aspects of dental morphology (e.g. curvature, complexity) or digestive physiology were more critical to dietary partitioning early in kangaroo evolution. Late Oligocene through middle Miocene dental macrowear and crown-height values amongst balbarine and stem macropodids (figs. 1C,D) generally suggest reliance upon low-abrasion foods. However, some middle Miocene samples from northwestern Queensland show that balbarines had more abrasive diets near the middle Miocene climatic optimum (fig 1C) which seems to capture an unsuccessful attempt by balbarine kangaroos to capitalize on more abrasive plant resources before their late Miocene extinction (*15*).

**Fig. 1.**
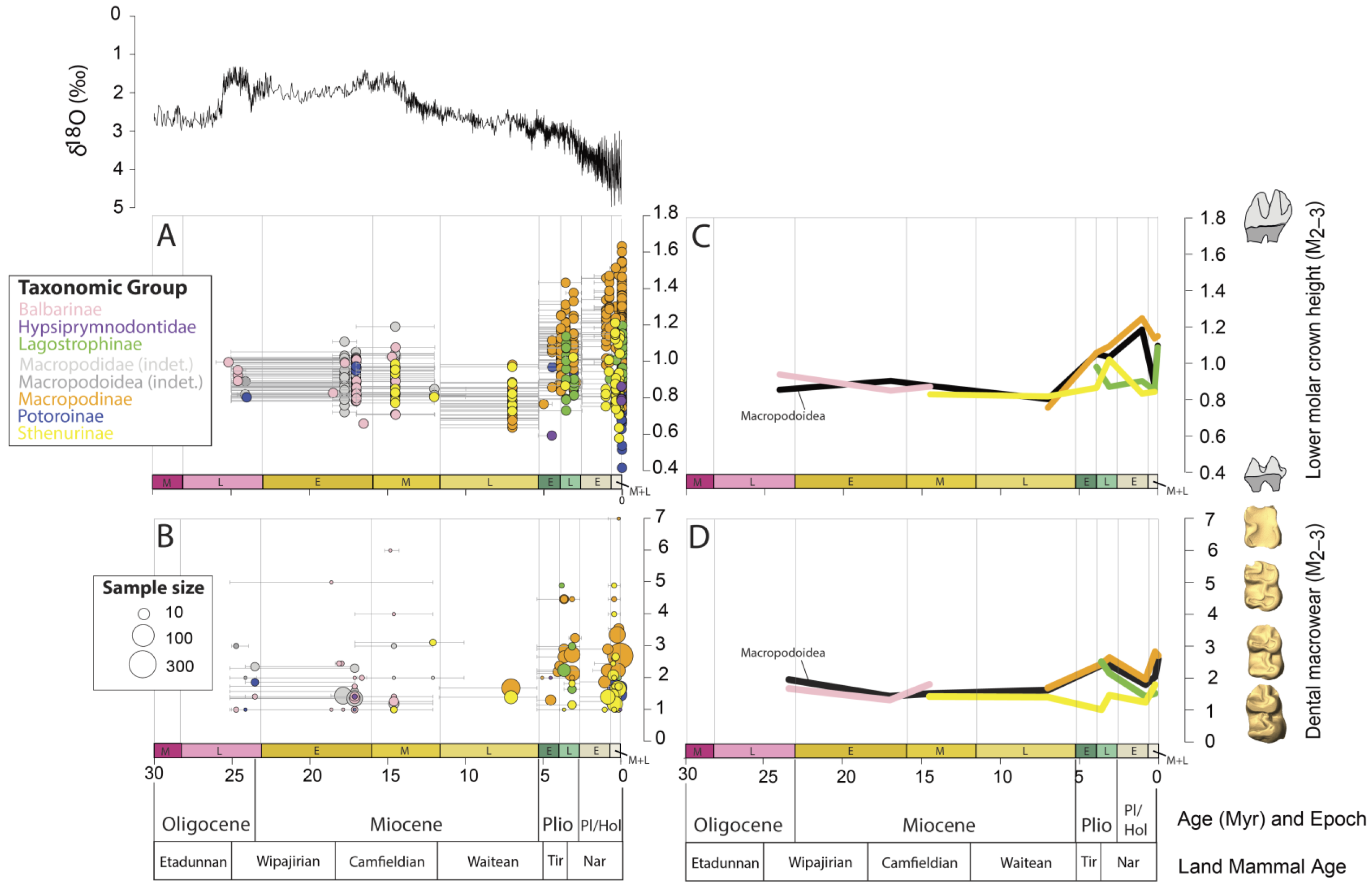
(**A**) Macropodoid molar crown-height (M_2 & 3_) and (**B**) dental macrowear (M_2, 3_) aligned with the global benthic foraminiferal oxygen isotopic curve (δ18O, black) (*31*). Median (**C**) macropodoid crown-height (M_2 & 3_) and (**D**) geometric mean macrowear (M_2, 3_) for macropodoid clades binned within geological sub-epochs. Land mammal ages follow (*32*). Abbreviations: Nar; Naracoortean, Tir; Tirarian, Pl; Pliocene, Hol; Holocene.

Comparative modelling reveals a middle Miocene split in adaptive strategies (fig 2A), with derived macropodids departing on a trajectory toward specialization as folivores, whereas hypsiprymnodontids and potoroines remained generalists or truffle consumers (fig 2A). Shifting between generalist and more specialized diets has been argued to require traversing of adaptive ‘valleys’ between fitness ‘peaks’ (*16*) and we hypothesize that transitional macropodid lineages exhibited faster taxonomic turnover, lower abundance and smaller geographic ranges than taxa closer to these trophic optima (e.g. *10, 17*).

**Fig. 2.**
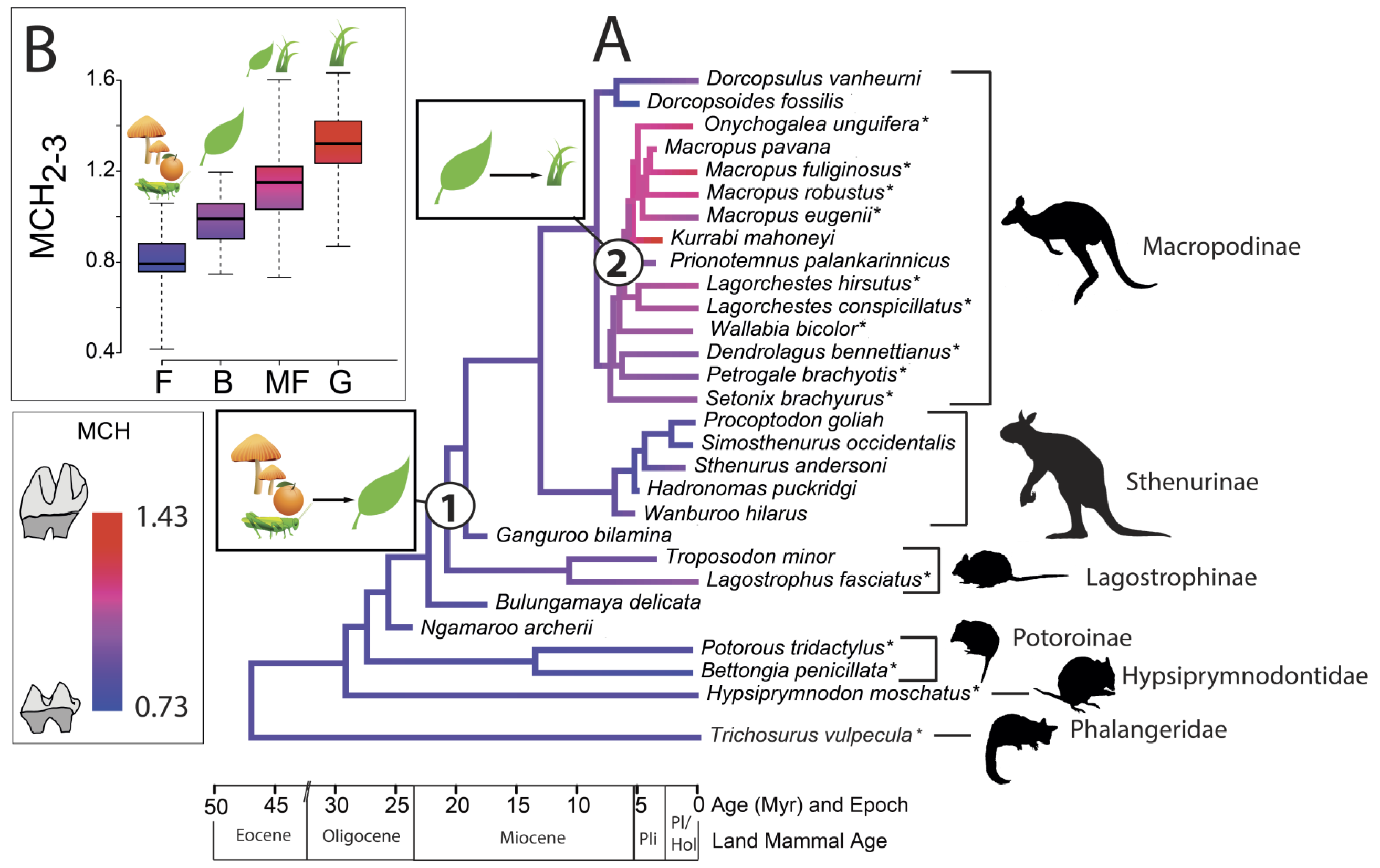
Phylogenetic reconstruction of crown-height in fossil and living macropodoids and relationship between diet and crown-height in living macropodoids. (**A)**. Phylogenetic reconstruction of molar crown-height evolution in crown-group Macropodoidea reveals an early Miocene diet shift from fungi to browse and an early Pliocene switch from browse to grass. (B) Extant grass-consuming macropodoids have significantly greater crown-heights than non-grass-consuming species. Diet abbreviations: ‘F’, Fungivore; ‘B’, Browser; ‘MF’, Mixed-feeder; ‘G’, Grazer.

Unexpectedly, we find no evidence for increased late Miocene macrowear or crown-height analogous to those interpreted as responses to aridity in Northern Hemisphere herbivores (*10, 11, 18*). Instead, late Miocene macropodids, represented by early members of the two most diverse kangaroo clades (Sthenurinae, Macropodinae), express even lower crown-height and macrowear levels than earlier macropodoids (figs. 1A, B). This is especially surprising given that central Australian faunal assemblages (e.g., Alcoota) should have been amongst the first to see grassland establishment (*19*). The low macrowear levels and low-crowned bilophodont molars typical of late Miocene macropodids, together with the basal phylogenetic position of modern browsing macropodids (e.g., *Dendrolagus*, *Dorcopsis, Setonix*) (*7–9*), indicate that browsing was ancestral for the subsequent sthenurine and macropodine radiations.

Marked crown-height and macrowear increases across the Miocene–Pliocene transition (fig.1) herald a major adaptive shift in kangaroo evolution. Molar crown-height increased by up to 40% in as little as 3 Myr (fig 1A), which is comparable to the fastest rates measured amongst Neogene ungulates (*10*). Large parameter estimates for selection pressure from the best-fitting phylogenetic comparative model (table S7) and the correlated increase of crown-height and macrowear suggest that accelerated dental evolution was driven by selection for improved dental durability. Although early Pliocene macrowear levels are somewhat higher than late Miocene levels (fig 1B), they are markedly lower than mid-and late Pliocene levels, suggesting that the rapid spread of late Pliocene grasslands intensified selective pressure for morphological diversification. Increasing molar crown-height was likely adaptive because it delayed loph collapse, the point at which bilophodont molars are relegated to a crushing rather than cutting modality (*20*). Supporting this, macrowear from the mid-Pliocene onwards is characterized by unprecedented loph destruction (fig 1B).

Average macrowear and crown-height levels are similar between Pliocene macropodins and non-macropodins (fig 3A); only during the late Pliocene do macropodins show evidence of high-wear diets (fig 3B). Based on the diet–crown-height relationship amongst extant kangaroos (Fig 2B), the highest-crowned mid to late Pliocene macropodins (Chinchilla, Bluff Downs) evidently consumed both grass and dicot leaves. This generalist diet fits with enamel δ13C values for species of *Macropus* and *Protemnodon* from Chinchilla (*21*). It suggests that the Pliocene was a period of trophic generalism amongst macropodines, with dietary specialization emerging surprisingly late, perhaps not until the early Pleistocene. This late arrival of specialized diets helps explain why most extant macropodines are mixed-feeders (*22*) and the rapid rates of dental evolution associated with generalist diets suggests less-commonly-consumed, fallback foods can be potent drivers of dietary adaptation (*23*).

**Fig. 3.**
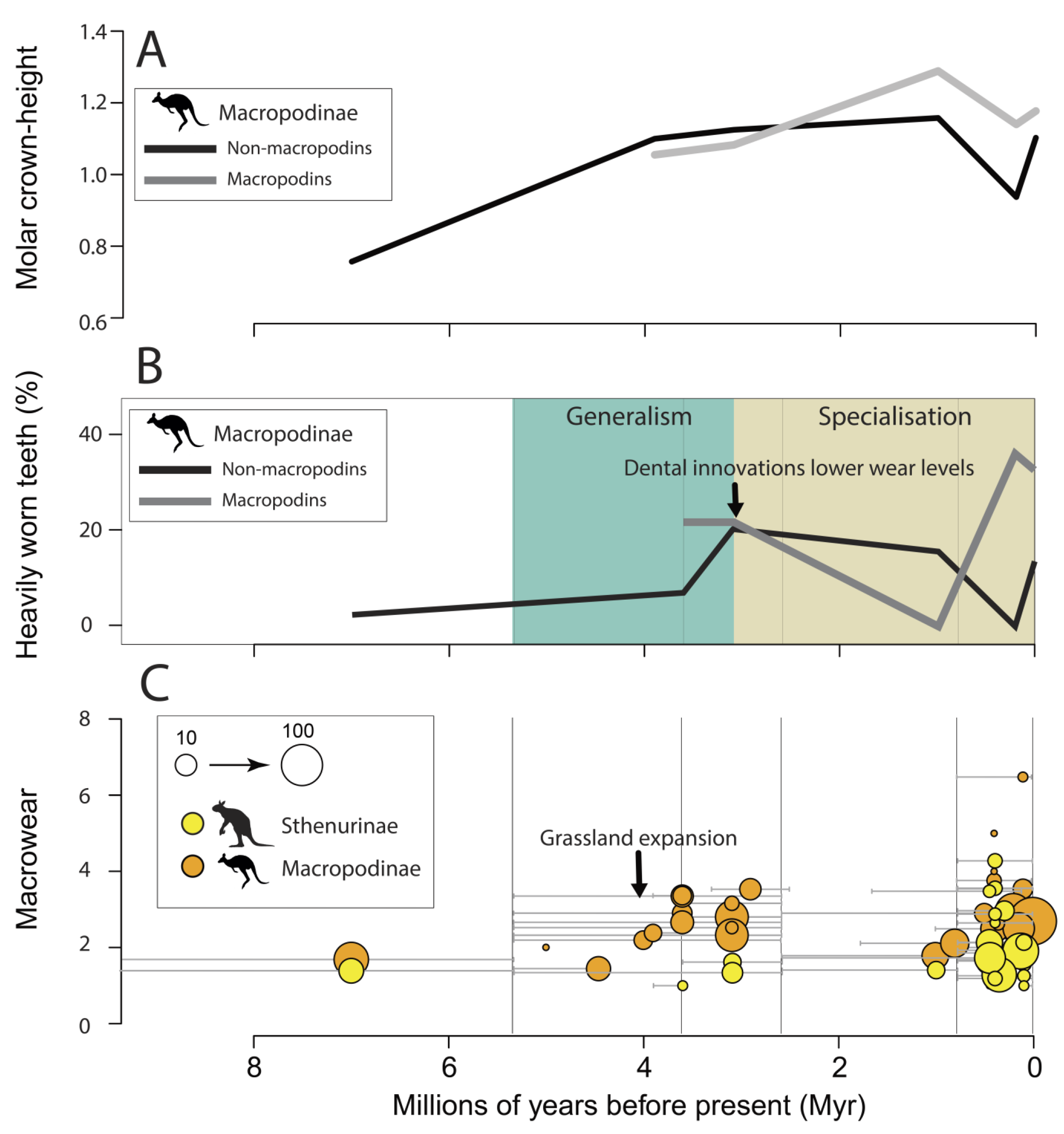
Dental macrowear evolution in macropodine and sthenurine kangaroos during the late Neogene. (**A**). Both Pliocene macropodins and non-macropodins show trends towards increasing molar crown–height; the extant pattern of high-crowned macropodins and lower-crowned macropodins emerged in the Pleistocene. (**B**). Early and middle Pliocene macropodin and non-macropodin kangaroos both consumed abrasive foods but late Pliocene dental innovations like high-crowned teeth enabled macropodins to specialize on grasses. **(C)**. Sthenurine and macropodine kangaroos differentiated into low and high abrasion diets respectively during the early Pliocene with sthenurines diversifying diets again during the middle and late Pleistocene.

The evidence for rapid macropodin dental evolution has important implications for the timing and context of kangaroo diversification. Most recent phylogenetic analyses support a short-fuse model (Fig 4A), where macropodin generic and many intrageneric splits are placed within the drying late Miocene (*7, 8*). However, this fits uncomfortably with our evidence for a Pliocene adaptive shift, as well as the absence of any known Miocene macropodins. Our data instead support the existence of two Pliocene events: 1) macropodin genera (Fig 4B) emerge during the early Pliocene ‘warm reversal’ (*24*); and 2) adaptively radiate during the arid late Pliocene and early Pleistocene as grassland expansion enables dietary partitioning.

**Fig. 4.**
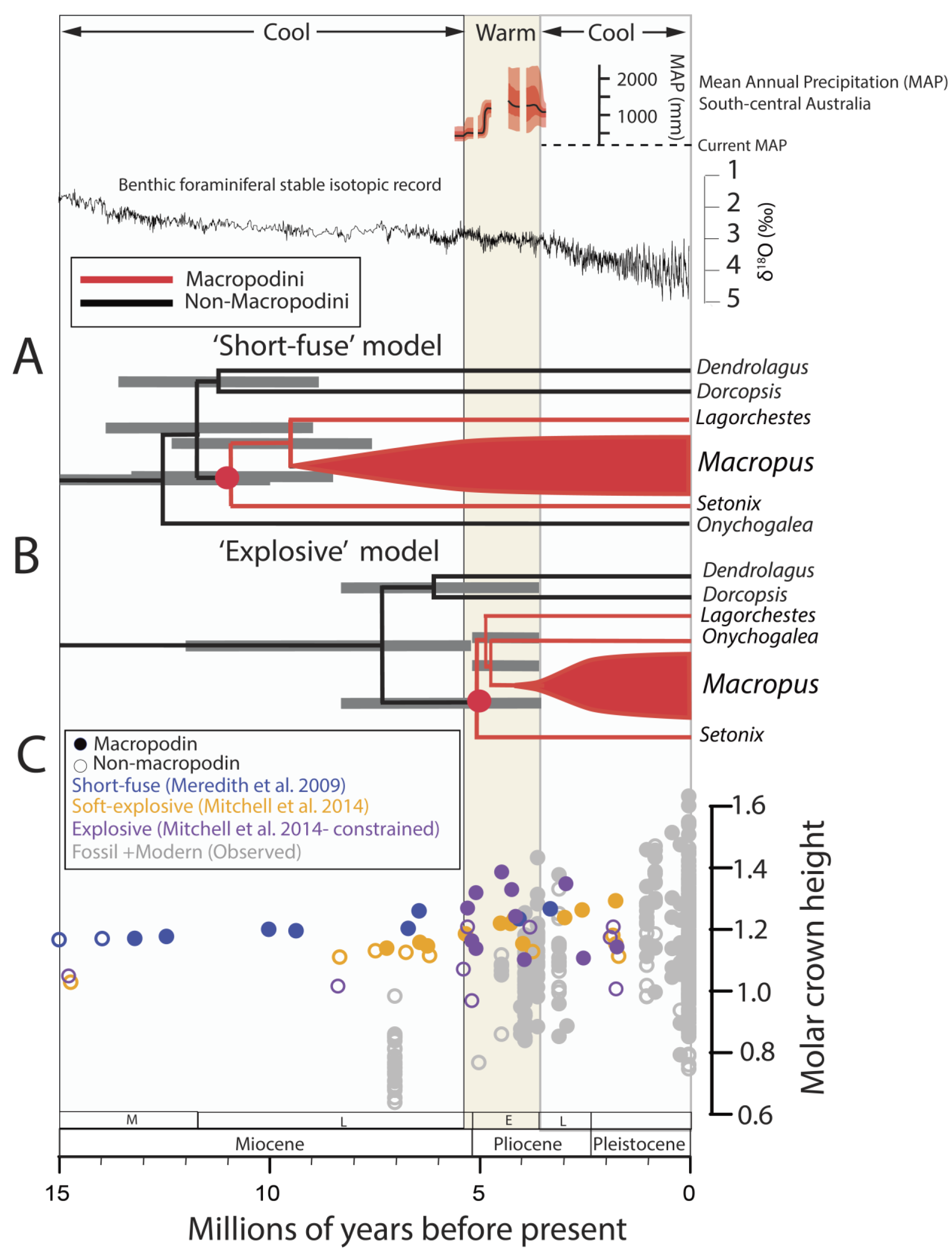
Alternative models of kangaroo diversification and climatic oscillations. (**A**). The ’short-fuse’ model links macropodin adaptive radiation with late Miocene aridification. **(B)**. An ‘explosive’ model implies diversification during the Early Pliocene warm-wet interval. Divergence times and error bars for the short-fuse model follow (*8*). Global stable oxygen isotopic record from (*31*) and Nullarbor Region (south-central Australia) mean annual precipitation from (*24*). (**C**). Restricting macropodin and dendrolagin diversification to the Early Pliocene is the only model which reproduces increased Pliocene crown-height.

This scenario implies accelerated molecular rates at the base of Macropodini. Under this model the occurrence of *Macropus* at 4.46 Myr (Hamilton) necessitates around four genus-level splits (*Macropus*, *Setonix*, *Lagorchestes*, *Onychogalea*) within less than 0.9 Myr. While rapid, this is still slower than speciation rates implied for some placental radiations (*4*). Or inference is supported by recent, whole-genome phylogenetic analyses that suggest rapid, even reticulated evolution within *Macropus* (*25*).

The limited Australian late Miocene fossil record means a hitherto-concealed Miocene macropodin radiation cannot be ruled out, but several lines of evidence favor a more rapid Pliocene diversification model. First, despite >50 years of collecting, the late Miocene Alcoota assemblage of central Australia has yielded no macropodins, but many specimens of the low-crowned dorcopsin *Dorcopsoides fossilis*, sthenurine *Hadronomas puckridgi*, and three as-yet-undescribed, low-crowned, non-macropodin kangaroos. Second, an extremely rapid basal divergence of macropodins would help explain the persistent difficulty in phylogenetically placing putative basal macropodins like *Setonix* and *Onychogalea* as well as evidence for substantial introgression at the base of Macropodini (*25*). Third, given grasslands were uncommon until the late Pliocene (*5*), an ‘explosive’ model would imply a much simpler scenario of dietary adaptation, with perhaps just a single acquisition of grazing, rather than as many as nine required by a short-fuse scenario (fig. S5). Finally, short-fuse trees invoke the existence of high-crowned macropodines 4–7 Myr before Pliocene grasslands and well outside the range of fossil crown-heights (fig 4C). Only by restricting the macropodin and dendrolagin radiations to the Pliocene do we recover the pronounced Pliocene increase in crown-heights captured by the fossil record (fig 4C). This raises the prospect that rock-wallabies and tree-kangaroos also did not originate until the early Pliocene.

The late Neogene has been interpreted as a phase of waning diversity amongst low-crowned browser groups (*18*), but we find that the Pleistocene diversification of sthenurine kangaroos (*26*) occurred almost entirely within a low-crowned region of morphospace (fig 1C). Sthenurine diversity more than doubles across the Pliocene–Pleistocene boundary (*26*) during a period when Australian terrestrial primary productivity was declining (*5, 24*). Sthenurine diversification thus contradicts expectations that intervals of high browser diversity are coupled to high primary productivity (*18*). In line with this peak in sthenurine species richness (*26*), the macrowear data reveal that sthenurine diets were diversifying (Fig 3C). The reliance of some sthenurines like *Procoptodon goliah* on chenopod shrubs, a dicot plant adapted to low-rainfall and high salinity (*27*), raises the prospect that expanded chenopod biomass around the middle Pleistocene arid shift (0.7 Myr) (*28*) could have been a key factor in this diversification. Evidence that sthenurine diversification was, if anything, gathering pace during the middle to late Pleistocene, and was closely coupled to dietary change through an interval of deepening aridity, discounts the possibility that such factors drove sthenurine extinction (*29*).

Aridity has been widely implicated in the diversification of modern clades (*6, 9*) but our data reveal a more dynamic picture where warm to cool oscillations promote taxonomic, and later ecological and morphological diversification. Warm–wet intervals are associated with diversification amongst other mammalian groups (*18, 30*), but a generalizable model of how these climatic perturbations are linked to diversification has yet to emerge. We propose that warm–wet conditions may ‘prime’ clades for rapid morphological and ecological diversification during ensuing arid intervals, perhaps by fostering trophic generalists that can later undergo bursts of speciation when ecological opportunity arrives (*16*). Future tests of this model which leverage the Cenozoic record of oscillating climate hold promise for revealing how climatically-driven ecological change drives adaptive diversification.

## Summary

Analysis of dental trait evolution shows that kangaroos rapidly diversified in response to Pliocene environmental change rather than Miocene aridification

## Acknowledgements

We thank the hundreds of volunteers, students and scientists who collected and prepared specimens used in this analysis. For specimen loans and access we thank Y.-Y. Zhen and S. Ingelby (Australian Museum), K. Spring and H. Janetski (Queensland Museum), A. Camens and L. Nink (Flinders University), D. Pickering and T. Ziegler (Museum Victoria), R. Palmer (Australian National Wildlife Collection), M.-A. Binnie and D. Stemmer (South Australian Museum), M. Siverson, H. Ryan, and L. Umbrello, (Western Australian Museum), A. Yates (Museum and Art Gallery of the Northern Territory) and P. Holroyd (University of California Museum of Paleontology). L. Hlusko and D. Polly are thanked for comments on an earlier version of this paper. R. Meredith kindly provided a copy of his divergence time tree. M. Rücklin generously supported completion of this paper while A. Couzens was a Naturalis postdoctoral fellow.

## Data materials and availability

All data, code, and materials used in the analysis are available on dryad.

## Supplementary Materials

Materials and Methods

Supplementary Text

